# Engineering Planar Gram-Negative Outer Membrane Mimics Using Bacterial Outer Membrane Vesicles

**DOI:** 10.1101/2023.12.11.570829

**Authors:** Aarshi N. Singh, Meishan Wu, Tiffany T. Ye, Angela C. Brown, Nathan J. Wittenberg

## Abstract

Antibiotic resistance is a major challenge in modern medicine. The unique double membrane structure of gram-negative bacteria limits the efficacy of many existing antibiotics and adds complexity to antibiotic development by limiting transport of antibiotics to the bacterial cytosol. New methods to mimic this barrier would enable high-throughput studies for antibiotic development. In this study, we introduce an innovative approach to modify outer membrane vesicles (OMVs) from *Aggregatibacter actinomycetemcomitans,* to generate planar supported lipid bilayer membranes. Our method first involves the incorporation of synthetic lipids into OMVs using a rapid freeze-thaw technique to form outer membrane hybrid vesicles (OM-Hybrids). Subsequently, these OM-Hybrids can spontaneously rupture when in contact with SiO_2_ surfaces to form a planar outer membrane supported bilayer (OM-SB). We assessed the formation of OM-Hybrids using dynamic light scattering and a fluorescence quenching assay. To analyze the formation of OM-SBs from OM-Hybrids we used quartz crystal microbalance with dissipation monitoring (QCM-D) and fluorescence recovery after photobleaching (FRAP). Additionally, we conducted assays to detect surface-associated DNA and proteins on OM-SBs. The interaction of an antimicrobial peptide, polymyxin B, with the OM-SBs was also assessed. These findings emphasize the capability of our platform to produce planar surfaces of bacterial outer membranes, which in turn, could function as a valuable tool for streamlining the development of antibiotics.

## Introduction

The escalating challenge of antibiotic resistance in modern healthcare necessitates urgent attention and innovative solutions to preserve the efficacy of antimicrobial treatment^1,2^. To address this pressing issue, there is a need to explore new methods for antibiotic development and testing. In light of the growing resistance to conventional antibiotics, antimicrobial peptides (AMPs) have gained prominence as an alternative^3–5^. They target bacterial membranes, reducing the potential for resistance to develop. However, evaluating the interaction of AMPs with bacterial membranes presents a considerable challenge, as it involves creating a membrane model system that mimics bacteria while enabling in vitro testing.

Gram-negative bacteria display distinct cellular architecture characterized by a double membrane structure consisting of inner and outer membranes^6^. These membranes encase a relatively thin layer of peptidoglycan. The outer membrane in gram-negative bacteria exhibits heterogeneity, with the outer leaflet comprising lipopolysaccharide (LPS) and outer membrane proteins, while the inner leaflet is predominantly composed of phospholipids^6^. For antibiotics, specifically AMPs, to be successful for gram-negative bacteria treatment, they must first interact with or be permeable through the outer membrane; therefore any changes to the outer membrane can lead to resistance^7,8^

Having a planar surface that mimics the bacterial outer membrane offers multiple advantages. For example, it provides a valuable tool for assessing interactions between potential antibiotics, such as AMPs, and the bacterial membrane using a variety of analytical techniques, such as surface-based biosensors atomic force microscopy^9,10^, fluorescence microscopy^11^, and quartz crystal microbalance^12^. In recent years, the utilization of synthetic lipids to create planar bilayers has emerged as a significant biotechnological advancement, and it has also been employed in the past for testing the effect of AMPs on gram-negative bacteria, where bacterial lipids with known composition were mixed to prepare vesicles^13–15^. However, synthetic lipid vesicles have several disadvantages, including that their composition is not an exact mimic of the outer membrane. In addition, these mimics do not contain proteins, which play a prevalent role in the outer membrane. An alternative to using synthetic lipids for vesicle production is to extract lipids directly from the bacteria^14,16,17^. However, a disadvantage of this method is that when bacterial lipids are extracted, there will be lipids present from both the outer and inner membranes of the bacteria, and the origin of the lipids cannot be precisely determined.

Tethered bilayer lipid membrane (tBLM) is another avenue that can be used prepare lipid bilayers to mimic the cell surface. In a tBLM system, a synthetic bilayer is formed on a tethered surface. This method allows for the used of various substrates, such as gold, which might not be feasible for bilayer formation using the vesicle rupture method^18,19^. Depending on the tethering approach, this setup allows for a cushion beneath the bilayer, effectively mimicking a cell surface, which allows for AMP insertion^20^. However, the challenges listed above associated with synthetic lipids remain, as the bilayer in this method is still constructed using synthetic lipids.

Gram-negative bacteria bud their outer membranes to generate nanoscale structures known as outer membrane vesicles (OMVs)^21,22^. These OMVs are crucial for bacterial communication and cargo transport. Furthermore, they play a significant role in pathogenicity as they can carry and deliver toxins to host cells^23–25^. Because OMVs originate from the bacterial outer membrane, they provide valuable insights into the composition of the outer membrane, including details about LPS, lipid composition, and proteins^26,27^. Given that OMVs closely mimic the outer membrane of their parent bacteria, they present a promising opportunity to serve as bacterial membrane mimics. Previously, Richter et al. introduced an assay based on OMVs, which enables the detection of antibiotic permeation, emphasizing the utility of outer membrane vesicles as a model for antibiotic development^28^. While this method excels in detecting antibiotics that permeate through the outer membrane, sensor-based applications were not demonstrated. In order to be used in many sensor-based applications a planar SLB is advantageous. However, unlike zwitterionic liposomes, OMVs do not spontaneously rupture on glass or silicon dioxide surfaces to form a planar supported lipid bilayer (SLB). This phenomenon can be attributed to the presence of anionic lipids, such as phosphatidylglycerol and LPS in OMVs, which hinders the electrostatic interactions between OMVs and glass necessary for rupture to occur.

Daniel and coworkers reported a method to make OMV-based supported lipid bilayers^29,30^. In this method, the OMVs were adsorbed in a high density on a surface, followed by the addition of synthetic liposomes to induce the rupture of OMVs. Recently, in addition to OMVs derived from Gram-negative bacteria, reports using vesicles derived from Gram-positive bacteria have also emerged, highlighting the versatility of this method^31^. While this method does allow for the formation of a surface incorporating the OMVs, it does have limitations, including that it requires the OMVs to be densely adsorbed, requiring a high concentration of OMVs. In addition, a high concentration of synthetic lipid was used to induce the rupture of OMVs, which dilutes the OMV components in the surface, and there was no direct measure of the how much synthetic lipid was present in the bilayer.

Our goal in this work was to create hybrid outer membrane vesicles (OM-Hybrids) that can spontaneously rupture on a silica dioxide surface to create outer membrane-supported bilayers (OM-SB). Previously, OMVs were shown to fuse with eukaryotic cells to form prokaryotic-eukaryotic hybrids for drug delivery using methods like sonication and extrusion^32–36^. Additionally, Pace et al. described a method involving cell lysis to create vesicles, followed by their fusion with synthetic liposomes using sonication, leading to their ability for spontaneous rupture and supported bilayer formation^37^. While this method was effective, our objective was to investigate the feasibility of using freeze-thaw (FT) as an alternative approach to create OM-Hybrids. Previously, rapid FT has been reported as a suitable technique for lipid-lipid mixing^38^. Moreover, FT has demonstrated success in generating exosome-liposome hybrids, which can be utilized for drug delivery^39,40^. FT has previously been reported to successfully load OMVs with molecular cargo^41–44^. This suggests that FT treatment effectively induces membrane disruption and reformation within OMVs, a property that can be harnessed for developing OM-Hybrids that can subsequently be used to form OM-SBs.

As a proof of concept, we utilized OMVs produced by *Aggregatibacter actinomycetemcomitans,* a Gram-negative bacterium associated with severe periodontal diseases^45^ that has been observed to be resistant to commonly used antibiotics^48,49^. As part of its virulence, *A. actinomycetemcomitans* produces leukotoxin (LtxA), which is secreted as a free protein and is also released in association with OMVs, where it resides on the surface along with DNA ^50–52^. This highlights the critical role of OMVs in pathogenesis, which is why we seek to utilize them, given their relevance in the disease process. For the synthesis of OM-Hybrids, we utilized liposomes consisting of phosphatidylcholine (POPC) and a small amount of PEGylated phosphatidylethanolamine (PEG-PE). The PEG moiety adds an extra cushion to the resultant OM-SB that prevents the bilayer from touching the surface, which prevents proteins from becoming pinned on the surface^37,53^.

We assessed the OM-Hybrid formation through techniques such as fluorescence quenching and dynamic light scattering (DLS), providing insights into their characteristics. We also determined the diffusion coefficient of lipids in the OM-SB through fluorescent recovery after photobleaching (FRAP) and observed that OM-SB exhibited a lower diffusion coefficient compared to the POPC/PEG-PE supported lipid bilayer (POPC/PEG-PE SLB). Additionally, we utilized quartz-crystal microbalance with dissipation monitoring (QCM-D) to assess OM-SB formation from OM-Hybrids. Finally, we investigated the presence of surface-associated DNA and proteins on the OM-SBs, demonstrating that our hybrid formation method preserves the integrity of the OMVs and their surface-associated components. Finally, we showed that an antimicrobial peptide, polymyxin B, aggregates on the surface of OM-SBs. This method exhibits versatility, facilitating the generation of planar membranes possessing bacterial biomolecules. Furthermore, its gentle nature ensures the preservation of critical outer membrane constituents.

## Experimental Section

### Reagents and chemicals

The following lipids were bought from Avanti polar lipids: 1-palmitoyl-2-oleoyl-glycero-3-phosphocholine (POPC), 1,2-dioleoyl-*sn*-glycero-3-phosphoethanolamine-N-[methoxy(polyethylene glycol)-5000] (ammonium salt) (PEG-PE). Octadecyl rhodamine B Chloride (R18), DiI, and DiO membrane labels were bought from Invitrogen. Triton X-100 was purchased from Sigma-Aldrich. Polydimethylsiloxane (PDMS) was purchased from Dow. YO-PRO-1 was purchased from Biotium. All experiments were performed in Tris buffer composed of 150 mM NaCl and 10 mM Tris-base (pH 7.0).

### Liposome preparation

A molar ratio of 99.5% POPC to 0.5% PEG-PE was dissolved in chloroform. The resulting solution was mixed within a glass vial, followed by the evaporation of the solvent to yield thin lipid films. The lipid films were rehydrated in Tris buffer with a final lipid concentration of 1 mg/mL, followed by gentle vortexing and bath sonication for 10 minutes. The resulting liposomes were extruded through a 50 nm pore diameter polycarbonate membrane (Cytiva Life Sciences) using a Mini-Extruder (Avanti Polar Lipids) with 23 passes.

### DiI/DiO incorporation in OMVs and liposomes

10 µM of DiI or DiO dissolved in DMSO was added to the either OMVs or liposomes. The solution was mixed by pipetting up and down followed incubating at 37 °C for 1 hour.

### Purification of A. actinomycetemcomitans OMVs

*A. actinomycetemcomitans* strain JP2 was raised in trypticase soy broth (30 g/L, BD Biosciences) and yeast extract (6 g/L, BD Biosciences) with the addition of 0.4% sodium bicarbonate (Fisher Scientific), 0.8% dextrose (BD Biosciences), 5 μg/mL vancomycin (Sigma-Aldrich), and 75 μg/mL bacitracin (Sigma-Aldrich). An initial culture was grown at 37 °C for 16 hours in a candle jar, then used to inoculate a larger culture that was allowed to grow for 24 hours at 37 °C. The bacteria were centrifuged once for 10 minutes at 10,000 × g, twice for 5 minutes at the same speed, and then filtered through a 0.45 μm filter. Following a 30-minute ultracentrifugation at 105,000 × g, the pellet containing the OMVs was resuspended in PBS (pH 7.4) and ultracentrifuged once more. Lastly, the final pellet was resuspended in PBS.

### OMV lipid concentration measurements

The lipid content of the OMVs was quantified by comparing the fluorescence intensity to a calibration curve generated with liposomes (POPC) at known concentrations. Both liposomes and OMVs were labeled with the FM 4-64 lipophilic dye (ThermoFisher). Fluorescence measurements were taken using a fluorometer with an excitation wavelength of 515 nm and an emission wavelength of 640 nm.

### OM-Hybrid preparation

Equal volumes of 0.2 mg/mL POPC/PEG-PE liposomes were mixed with OMVs that had a total lipid concentration of 4.5 mM. The solution was rapidly frozen at -196 ℃ using liquid nitrogen and subsequently thawed at 45 ℃ using a hot water bath. The FT cycle was repeated ten times, and at the end of the tenth cycle, the resultant OM-Hybrid vesicles were left in the water bath for 10 minutes.

### Dynamic light scattering (DLS)

DLS analysis was used to measure the diameters of liposomes, OM-Hybrids, and OMVs. Both OMVs alone and liposomes alone underwent ten FT cycles to make a direct comparison with OM-Hybrid production. Liposomes were diluted in Tris buffer to a final concentration of 0.25 mg/mL and OMVs and OM-Hybrids were diluted in Tris buffer with a dilution factor of 1:100. We used an ALV/CGS-3 Compact Goniometer System to make DLS measurements. At a wavelength of 623.8 nm and a scattering angle of 90°, data were gathered for 120 seconds in triplicate. The ALV software used a number-weighted regularized fit with an allowed membrane thickness (r*) of 5 nm to calculate the size distribution.

### Zeta potential measurement

OMVs, OM-Hybrids, and POPC-PEG/PE liposomes were subjected to zeta potential measurements. OMVs and OM-Hybrids were diluted 1:10 in sterile milliQ water to minimize salt presence. POPC-PEG/PE liposomes were diluted to 0.2 mg/mL in milliQ water as well. Zeta potential was measured at 25°C using a Zetasizer Nano ZS (Malvern).

### Lipid mixing study

0.2 mg/mL of POPC/PEG-PE liposomes containing 10 mole % R18 were prepared and mixed with non-fluorescent OMVs (4.5 mM lipid concentration) in equal volume, followed by rapid freeze thaw, as described above. The samples were then diluted 1:101 and added to quartz cuvette for fluorescence spectroscopy. Fluorescence emission spectra were collected using a Fluorolog TCSPC fluorometer (Horiba Scientific). The excitation was held constant at 500 nm while the emission was collected from 530 – 700 nm.

### Fluorescence recovery after photobleaching

Glass coverslips were cleaned first with isopropyl alcohol, then submerged in an aqueous 2% sodium dodecyl sulfate (SDS) solution for at least 30 minutes. The glass coverslips were thoroughly rinsed with milliQ water and then exposed to UV-Ozone (UV/Ozone ProCleaner Plus, BioForce Nanosciences) treatment for a duration of 10 minutes. To create small wells, first an ultra-flat PDMS slab was prepared, and the well was created within the PDMS slab using a 6 mm biopsy hole punch. The PDMS well was plasma treated for 90 seconds and then attached to the clean glass coverslip. OM-Hybrids labeled with DiI or 0.2 mg/mL of POPC/PEG-PE liposomes labeled with DiI were incubated on the glass coverslip for 1 hour and, then the resulting planar bilayers were washed with Tris buffer. The bilayers were imaged using an inverted microscope (Eclipse Ti, Nikon) equipped with a 100× oil immersion objective with a 1.49 numerical aperture. Fluorescence was excited using an LED light engine (Aura II, Lumencor) and appropriate filter cubes. Images were captured with a 2048 × 2048 pixel sCMOS camera (Orca Flash 4.0 v2, Hamamatsu). The planar bilayers were photobleached using a 405 nm laser (50 mW) pulse for 1 second and fluorescence recovery was captured at 1 second intervals for 60 seconds.

### Quantifying POPC-PEG in present in OM-SB

DiO-labeled POPC/PEG-PE liposomes were prepared and underwent OM-Hybrid formation by mixing them with DiI-labeled OMVs and FT cycles. These OM-Hybrids were used to create an OM-SB. Additionally, a separate equivalent volume and concentration of DiO-labeled liposomes used in OM-SB underwent FT cycles and were then exposed to clean glass to form a POPC/PEG-PE SLB. Subsequently, the fluorescence intensity from the OM-SB, POPC/PEG-PE, and OMVs (as a noise control) in both the DiI and DiO channels was compared. To calculate the percent of POPC/PEG-PE (labeled with DiO) present in OM-SB we used the formula below:

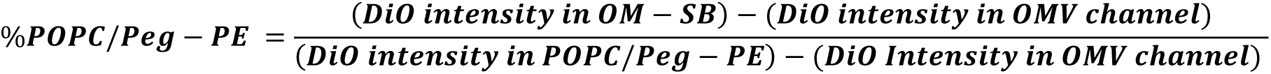

### Quartz crystal microbalance

A QSense Explorer E1 QCM-D instrument from Biolin Scientific, equipped with a 5 MHz AT-cut SiO_2_-coated quartz crystal (Nanoscience Instruments), served as the platform for conducting QCM-D studies. In the flow-cell, OM-SB hybrids or liposomes at a concentration of 0.20 mg/mL, or OMVs with a lipid concentration of 2.45 mM were introduced, flowing at a rate of 100 µL/min. While the 1st, 3rd, 5th, 7th, 9th, and 11th overtones were monitored, only the data corresponding to the 3rd harmonic is reported. The flow-cell temperature was held constant at 23 ℃.

### OM-SB DNA assay

OM-SB and POPC/PEG-PE bilayers were formed on clean glass coverslips as described above. After removing excess vesicles, the PDMS well was filled with a 1:1000 dilution of YO-PRO-1 (Biotium), which was then incubated for an hour. Bilayers were observed using an inverted microscope (Nikon Eclipse Ti) fitted with a 100× oil immersion objective with a 1.49 numerical aperture after the extra dye had been removed. A FITC (Chroma), and an LED light engine (Aura II, Lumencor) were utilized to excite and observe the fluorescence. A 2048 × 2048 pixel sCMOS camera (Orca Flash 4.0 v2, Hamamatsu) was used to collect images. ImageJ (NIH) was used to analyze and overlay images.

#### Protein labeling on OM-SB

OM-SB and POPC/PEG-PE bilayers were formed as described above. After washing the bilayers, 1 µg/mL of NHS Atto 488 was prepared in Tris buffer supplemented with 0.2 mM MgCl_2_, followed by 1 hour incubation. The NHS Atto 488 was thoroughly washed, and the bilayers were imaged using the parameters described in the section above. The mean intensity calculations were performed using ImageJ (NIH), over 10 individual images.

#### FITC - polymyxin B interaction with OM-SB

OM-SB and POPC/PEG-PE bilayers were formed as described above. After washing the bilayers, 1:1000 dilution of 1.25 mM FITC-polymyxin B stock was added to the chamber, followed by 1 hour incubation. The bilayer and the polymyxin B were imaged as described in previous sections. The 3D surface intensity plot was prepared using ImageJ (NIH).

#### Statistical analysis

All statistical analyses were performed using GraphPad prism V 10.0.3.

## Results and Discussions

### Formation of OM-Hybrids

The establishment of planar SLBs through the rupture of synthetic liposomes on glass and SiO_2_ surfaces is a common technique widely used for the creation of planar bilayer surfaces^54^. In general, liposomes composed of zwitterionic phospholipids, such as POPC, readily adsorb and rupture to form SLBs on glass and SiO_2_ surfaces. OMVs released by gram-negative bacteria exhibit the opposite behavior—they do not rupture upon contact with glass surfaces; instead, they sparsely adhere to the glass and remain intact **(Figure S1)**. In order to surmount this challenge, our aim was to engineer hybrid vesicles that would be amenable to planarization by vesicle rupture. DiI labeled OMVs were mixed with non-fluorescent liposomes (99.5% POPC, 0.5% PEG-PE) in an equal volume. The mixture was rapidly frozen using liquid nitrogen, followed by thawing at 45°C. This process was repeated 10 times to ensure a complete lipid mixing **(Figure 1).** After the completion of the tenth cycle, the OM-Hybrids were immersed in a warm water bath at 45°C and subsequently exposed to a clean glass coverslip to form a planar OM-SB.

**Figure 1:**
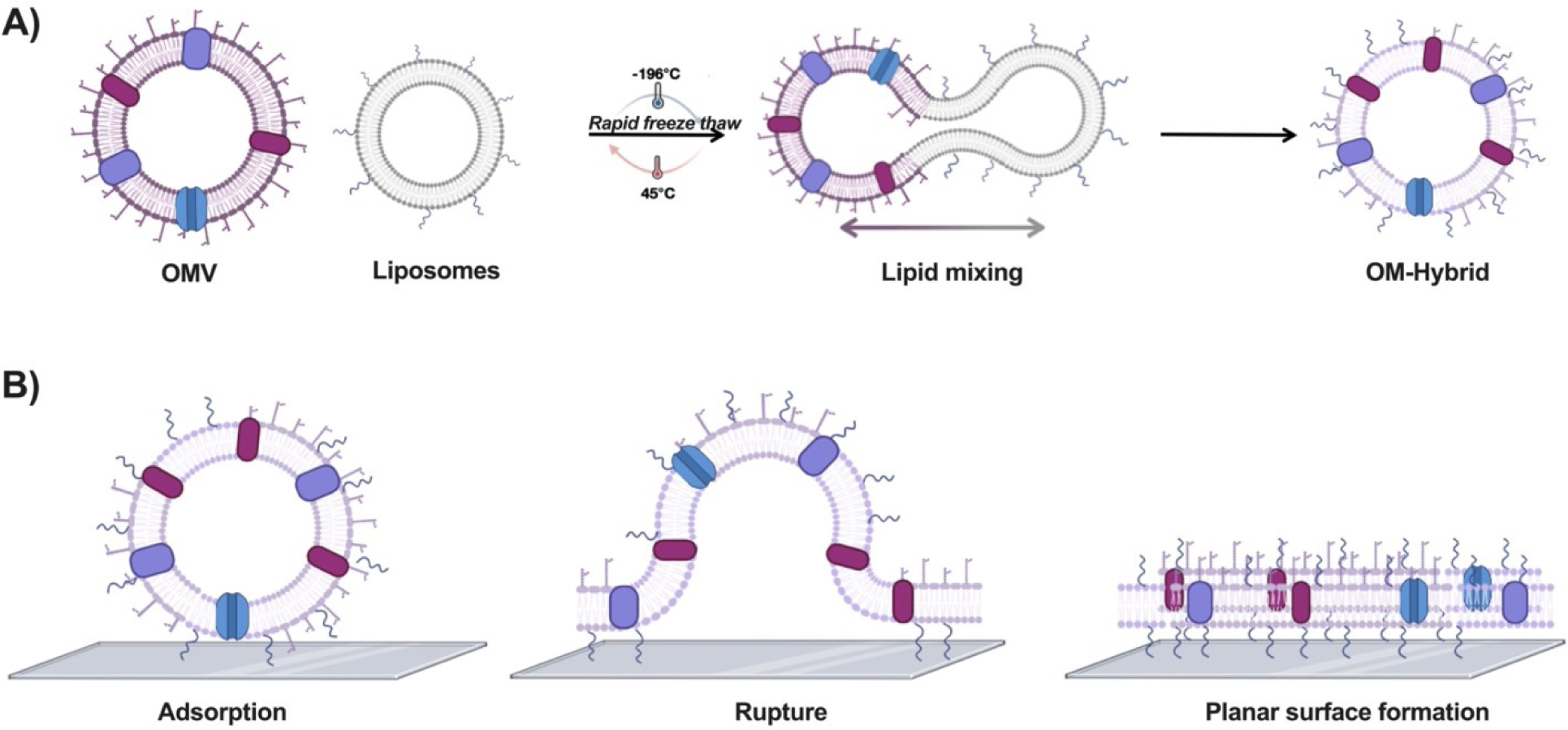
Schematic representation of the process: **(A)** Equal volumes of OMVs and POPC/PEG-PE liposomes are mixed and subjected to rapid FT treatment. The FT treatment induced lipid breaking and reformation, which results in lipid mixing and formation of OM-Hybrids. **(B)** OM-Hybrids adhere to the clean glass surface, initiating interactions that lead to their rupture and the formation of a planar surface.

### Lipid mixing between POPC/PEG-PE and A. actinomycetemcomitans OMVs

To investigate the interaction between liposomes and OMVs and the formation of OM-Hybrids, we employed the R18 fluorescent dye. This dye is recognized for its self-quenching behavior at elevated concentrations and has previously been demonstrated as effective for studying OMV-OMV fusion^55^. The experimental strategy involved creating POPC/PEG-PE liposomes that contained 10 mole % of R18, and subsequently monitoring the fluorescence enhancement upon FT treatment with non-fluorescent OMVs. If lipid mixing was occurring, an increase in fluorescence was expected due to R18 dilution in the OM-Hybrids. Initially, as a validation of the concept, we exposed R18-labeled POPC/PEG-PE liposomes to a treatment with 0.1% Triton. Triton lyses the liposomes, liberating the R18 molecules and resulting in an increase in observed fluorescence **(Figure S2).** We noted a significant 6.3-fold enhancement in dye emission following the complete lysis of liposomes and the subsequent release of the R18 fluorescent dye.

Equal volumes of R18 labeled POPC/PEG-PE liposomes and OMVs were mixed and subjected to either five or ten FT cycles. After subjecting the POPC/PEG-PE liposomes and OMVs to five FT cycles, we observed a modest 1.2-fold increase in fluorescence compared to FT POPC/PEG-PE liposomes alone **(Figure 2)**. However, upon performing ten FT cycles, we noted a statistically significant 1.5-fold increase in fluorescence. Consequently, we chose to implement ten FT cycles in our protocol. Furthermore, we sought to determine if the increase in fluorescence was a result of FT treatment. To investigate this, equal volumes of R18 labeled POPC/PEG-PE liposomes with non-labeled OMV were mixed, but were examined without the FT treatment. In this case, we observed a non-significant change in fluorescence intensity between liposomes and the non-treated samples **(Figure S3).** We also investigated if the R18 liposomes themselves are affected by the FT treatment. An insignificant change was observed, indicating that OMV-liposome fusion is the result of an increase in intensity **(Figure S4)**. This further underscores the essential role of the freeze thaw method in facilitating lipid mixing and OM-Hybrid formation.

**Figure 2:**
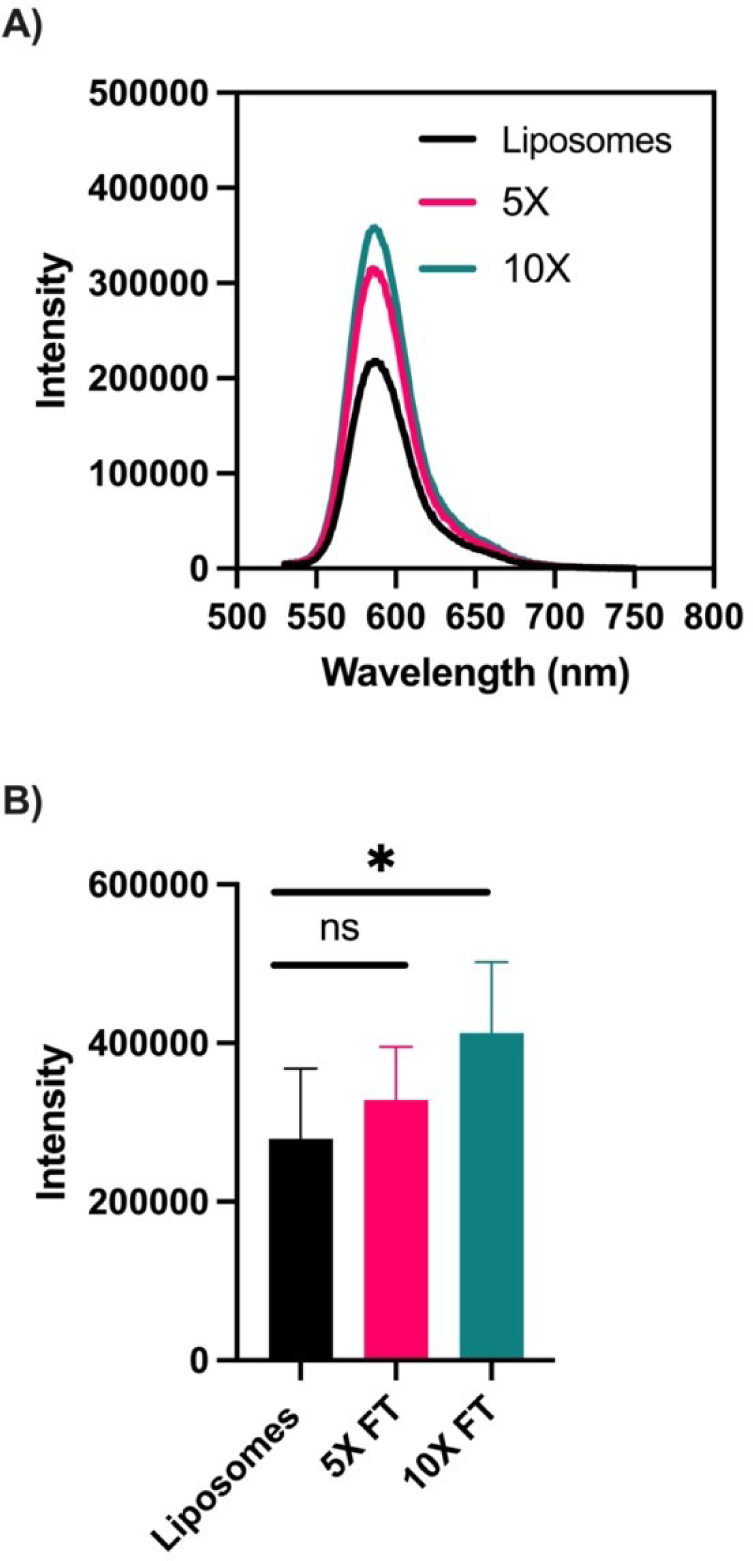
Quantifying lipid mixing through monitoring R18 fluorescence increase: **(A)** Representative spectra of the fluorescence intensity of R18 in liposomes alone (dark blue), OMV-Hybrids mixture subjected to five FT cycles, or ten FT-cycles (light blue and purple, respectively). **(B)** A non-significant increase in fluorescence intensity was observed after 5 FT-cycles (n = 5). However, a significant increase was observed after 10 FT-cycles (n = 5, p = 0.0454).

### Size and zeta potential analysis of OM-Hybrids

We used DLS to measure the diameters of OM-Hybrids. Each individual component of the hybrid, namely the OMV and the POPC/PEG-PE liposomes, underwent a FT process to ensure accurate evaluation of the treatment’s effects. Notably, we observed that the diameter of the OM-Hybrid was 102.3 ± 1.6 nm (mean ± s.d.), while the POPC/PEG-PE liposomes were 70.2 ± 6.9 nm and the OMVs were 140.8 ± 6.7 nm **(Figure 3)**. The mean diameter of the OM-Hybrids is in between the mean diameters of each component, indicating that OM-Hybrids are a mixture of these two components. This reinforces the notion that OM-hybrids exhibit properties akin to POPC/PEG-PE liposomes while retaining characteristics of OMV, confirming the occurrence of hybrid fusion.

**Figure 3:**
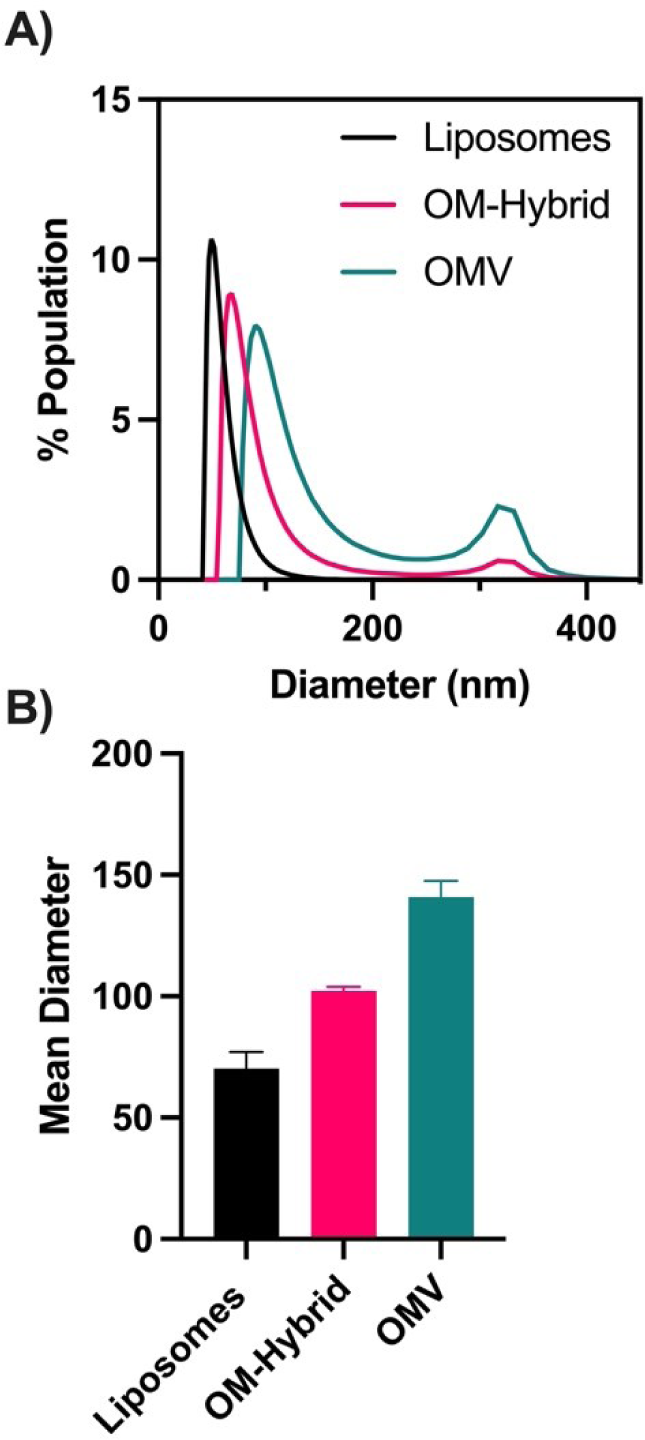
Size of OM-Hybrids and their individual components: **(A)** Representative diameter distributions of liposomes, OM-Hybrids, and OMVs (shown in blue, purple, and pink, respectively). **(B)** Quantification of the mean diameter of OM-Hybrids and their individual components (n=3).

To explore the impact of POPC-PEG/PE lipid fusion on the surface charge of OMVs, we conducted zeta potential measurements on OM-Hybrids and each individual component **(Figure S5)**. As anticipated, we observed that the outer membrane vesicles (OMVs) exhibited a higher negative potential compared to POPC/PEG-PE liposomes. There was no significant difference in the zeta potential between OM-Hybrids and OMVs. During the preparation of OM-Hybrids, the lipid content of bacterial OMVs is 4.5 mM, whereas the concentration of POPC/PEG-PE liposomes is only 0.026 mM, which is significantly lower compared to the amount of bacterial lipid present.

### OM-SB fluorescence microscopy and FRAP

A key feature of OM-Hybrids is their ability to spontaneously rupture on glass, forming an OM-SB. Membrane fluidity stands as a crucial attribute in biological systems, facilitating pivotal processes such as protein-lipid interactions, molecular transport, and the maintenance of cell membrane integrity. To verify the fluid nature of the OM-SB, our goal was to determine the lipid diffusion coefficients of OM-SB and POPC/PEG-PE SLB by analyzing their respective FRAP profile. OM-Hybrids were prepared with OMVs labeled with DiI, along with unlabeled POPC/PEG-PE, as described in the above section. To ensure a fair comparison of diffusion coefficients between OM-SB and POPC/PEG-PE SLB, we labeled the POPC/PEG-PE liposomes and subjected them to FT cycles at a consistent concentration of 0.2 mg/mL, mirroring the lipid concentration found in the OM-Hybrids. OM-Hybrids and POPC/PEG-PE liposomes were ruptured on clean glass surfaces, and the lateral diffusion of their lipids was examined using FRAP **(Figure 4A-B)**. We observed that the OM-SB exhibited a slower fluorescence recovery when compared to the POPC/PEG-PE SLB indicating that diffusion of components in the OM-SB is slower. Furthermore, it was noted that OM-SB exhibited a 76% recovery, while the POPC/PEG-PE SLB showed an 86% recovery of **(Figure 4C)**. We hypothesized that the presence of the biological factors such as proteins, porins, and LPS leads to a slower recovery in OM-SB. To test this, we varied the concentration of the synthetic lipid component in the formation of OM-SB, which led to a reduced amount of the OM component. We then observed their FRAP profiles **(Figure S6)** and noticed that with an increase in the synthetic lipid concentration, the OM-SB frap curves were similar to the POPC-PEG pure synthetic lipid curve. We reason that with additional lipid, the concentration of proteins and porins decrease, which leads to an increase fluorescent lipid mobility.

**Figure 4:**
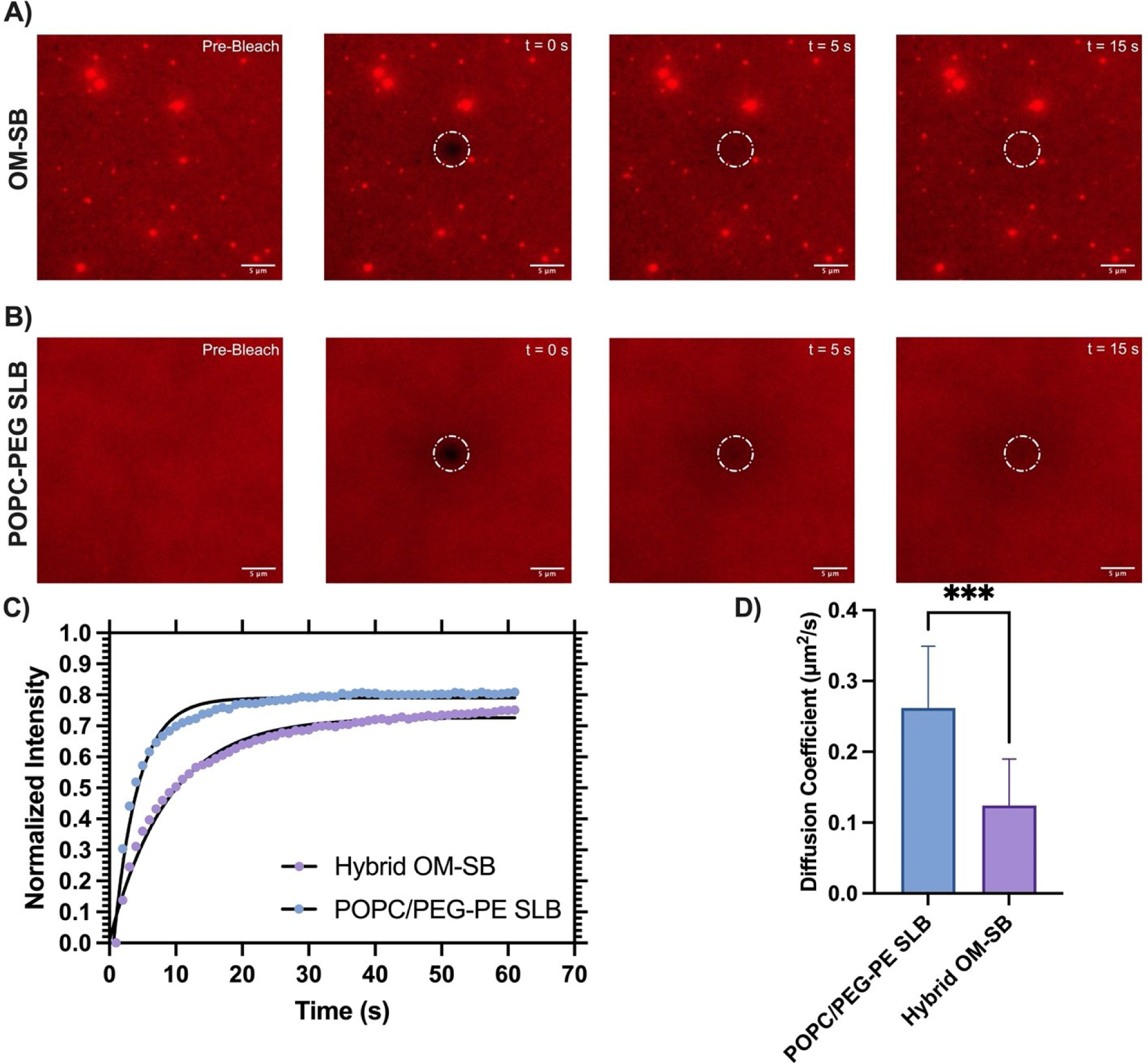
FRAP of OM-SB and POPC/PEG-PE SLB: **(A)** Fluorescence images of OM-SB **(B)** POPC/PEG-PE SLB during the bleach and recovery stages. The white dashed circle represents the bleach spot. Scale bar in **A** and **B** = 5 µm **(C)** Average FRAP recovery curves of OM-SB and POPC/PEG-PE SLB (n =12). The black lines represent the exponential fit used for analyzing the FRAP curves. **(D)** Diffusion coefficient of the OM-SB and POPC/PEG-PE SLB. A significant decrease in diffusion of OM-SB was observed (p = 0.0002; n=12).

To determine the diffusion coefficients for OM-SB and POPC/PEG-PE SLB, we fit the fluorescence recovery data to the single exponential equation, where *F_o_* is the *f*(t) value (fluorescence intensity) when *t* (time) is zero, plateau is the maximum recovery, and K is the rate constant which is expressed 1/s^56,57^. Using this fitted equation, we can solve for half-time for recovery (*t_1/2_*) where 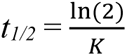.

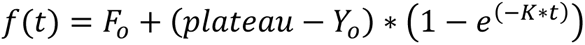

By obtaining the *t_1/2_* values from the FRAP curves fits, we were able to calculate the diffusion coefficients for these components using the equation below^58^.

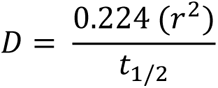

We observed that POPC/PEG-PE SLB had a diffusion coefficient of 0.26 ± 0.09 µm^2^/s while OM-SB had a diffusion coefficient of 0.12 ± 0.06 µm^2^/s **(Figure 4D)**. We hypothesize that the lower diffusion coefficient of the OM-SB suggests the presence of biological materials such as proteins and LPS within the bilayer that likely contribute to the reduction in diffusivity of the fluorescent lipids. It is worth noting that these values align with previously reported literature findings. Previously, it was reported that R18 fluorophore in POPC/PEG-PE had a diffusion coefficient of 0.28 µm^2^/s, while the R18 fluorophore in DOPC/PEG-PE had a diffusivity of 0.5 µm^2^/s ^29,30^. Studies by Daniel and coworkers on other planar bacterial membrane formation methods, have reported that the diffusivity of lipids in the bacterial outer membrane surface ranged from 0.33 to 0.4 µm^2^/s depending on the bacterial source of OMVs^29,30^. While these values are higher than those we have observed here, a likely reason for the discrepancy is the use of less of the synthetic lipid component in our method. Since our method uses only 0.2 mg/mL synthetic lipid, we see a lower diffusion, indicating a larger fraction of the membrane is of bacterial origin.

We conducted a series of experiments to explore the effectiveness of the FT method alone, without synthetic components, in forming a bilayer and causing OMV rupture. After subjecting a comparable concentration of OMVs to FT-treatment and subsequently adding them to a glass surface, our results revealed that freeze thaw OMVs exhibited adsorption but did not rupture on the glass surface **(Figure S7)**. This underscores the necessity of synthetic lipid integration for bilayer formation.

Next, we aimed to determine the minimum concentration of POPC/PEG-PE required to create OM-Hybrids that would rupture effectively. To minimize the use of POPC/PEG-PE liposomes, we tested various concentrations, mixing them with OMVs in equal volumes while varying the liposome concentration. Notably, we observed that 0.1 mg/mL did not result in OM-SB formation, producing only adsorbed vesicles. While both 0.5 mg/mL and 1.0 mg/mL liposomes produced OM-SBs, we found that 0.2 mg/mL was the lowest concentration that allowed us to maintain a substantial amount of the bacterial outer membrane component **(Figure S8)**. Interestingly, we also observed a reduction in bilayer fluorescence intensity as a function of increasing POPC/PEG-PE vesicle concentration **(Figure S8 E-H)**. In our bilayer formation experimental set-up, the fluorophore is exclusively present in the OMV, while the liposomes lack any fluorescent lipid. As the synthetic lipid concentration increases during OM-Hybrid and subsequent OM-SB formation, the fluorescence decreases, signifying a dilution of the OMV component.

Furthermore, we assessed the necessity of the FT process for OM-SB formation by conducting a negative control experiment in which OMVs and POPC/PEG-PE liposomes were mixed in equal volumes but were not subjected to FT treatment. In this case, we did not observe the formation of an OM-SB, confirming that the FT treatment is indeed essential for successful OM-Hybrid and OM-SB formation **(Figure S9).**

We also quantified the amount of POPC/PEG-PE in the OM-SB. DiO-labeled POPC/PEG-PE liposomes were prepared, mixed with DiI-labeled OMVs, subjected to FT cycles and exposed to glass surfaces. This process resulted in the formation of OM-SB, which contained both fluorophores **(Figure S10)**. We also prepared a SLB composed solely of POPC/PEG-PE as well as surface with adsorbed OMVs. While we observed similar fluorescence intensities between the OMVs and the OM-SB, there was a discernible reduction in the fluorescence intensity of the POPE/PEG-PE within the OM-SB compared to the POPC/PEG-PE SLB. We conducted additional quantification of the POPC/PEG-PE incorporation in both the OM-SB and POPC/PEG-PE SLB, revealing a significant decrease in DiO intensity in the former. By our calculations we found that approximately 32% of the OM-SB composition is derived from the POPC/PEG-PE liposomes.

### Real-time observation of OM-SB formation

Lastly, we tested the efficacy of the OMV and POPC/PEG-PE fusion. To do so, we observed real-time formation of OM-SB under TIRF microscopy **(Movie S1).** We observed that there were some vesicles that did not rupture on the glass. This implies that the formation of OM-Hybrids is not achieved by all OMVs. This finding aligns with the DLS results shown in **Figure 3**, which illustrate the continued presence of a segment of the larger OMV population within the hybrid structure.

### QCM-D analysis of OM-SB formation

To examine the formation pathway and biophysical properties of the OM-SB and POPC/PEG-PE SLB, we used QCM-D with sensor chips coated with SiO_2_. QCM-D is commonly used to analyze the frequency and dissipation properties of a surface^12^. In QCM-D measurements, a negative frequency shift is indicative of mass adsorption. Additionally, by monitoring dissipation, we gain insights into surface viscoelasticity. For instance, when liposomes or OM-Hybrids first adsorb intact to the surface, an increase in dissipation occurs because these structures can dissipate the vibrational energy of the quartz crystal sensor. However, as these vesicles rupture, a decrease in dissipation is observed, reflecting the planar bilayer’s rigidity and inability to dissipate as much energy.

Initially, we conducted an evaluation of the frequency and dissipation characteristics of OMVs adsorbing on SiO_2_-coated QCM-D sensors. We observed a final frequency shift of approximately -2.51 Hz and a dissipation of 1.33 × 10^-6^ **(Figure S11)**. It is noteworthy that OMVs containing LPS tend to exhibit lower frequencies, compared to liposome adsorption, when interacting with the QCM sensor, as LPS can interfere with mass detection, as previously observed with OMVs of *P. aeruginosa* and *E. cloacae*^30,59^. Next, we sought to study the rupture kinetics of POPC/PEG-PE vesicles and OM-Hybrids. We noticed that POPC/PEG-PE liposomes and OM-Hybrids had distinct rupture curves, indicating there are differences in the properties of the planar membranes formed from them **(Figure 5)**. Both types of vesicles displayed a comparable two-step rupture process, initially accumulating and then rupturing. However, the final frequencies were different, with OM-SB at -39.5 ± 0.7 Hz and POPC/PEG-PE SLB at -30.3 ± 2.4 Hz **(Figure 5A)**. OM-SB had a larger frequency change compared to POPC/PEG-PE, likely due to the presence of additional components (LPS, proteins, DNA, etc.) associated with the outer membrane of bacteria. Furthermore, our observations indicate that OM-SB exhibits higher viscoelasticity, as evidenced by a dissipation value of (3.7 ± 0.098) × 10^-6^, in contrast to POPC/PEG-PE SLB, which has a value of (1.8 ± 0.79) × 10^-6^ **(Figure 5B)**. POPC bilayer, without the addition of PEG-PE component has a reported dissipation around 0.55 × 10^-6^ and a frequency shift of around -28.3 Hz^60,61^. While the frequency characteristics of the POPC/PEG-PE SLB align with those of a POPC bilayer, we observe an increase in viscoelasticity in the POPC/PEG-PE SLB. We hypothesize that incorporating PEG-PE results in water association with PEG, leading to an increase in dissipation. A POPC-PEG/PE bilayer has been reported to have a frequency of ∼ -23 Hz and dissipation of ∼2.5 × 10^-6^ ^30^. These values are similar to the values that we reported.

**Figure 5:**
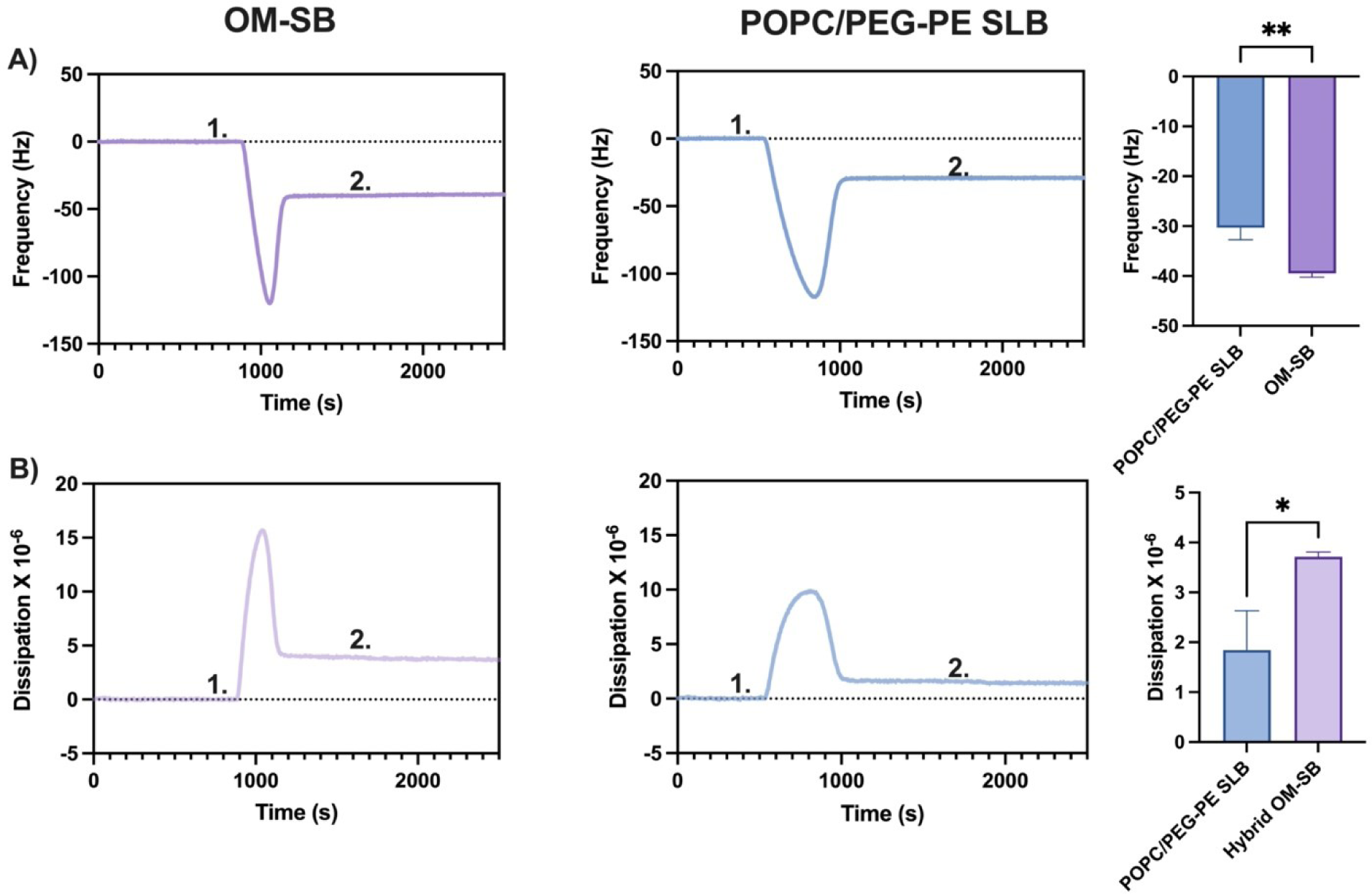
QCM-D of OM-SB and POPC/PEG-PE SLB: **(A)** Frequency analysis of OM-SB (shown in purple) and POPC/PEG-PE SLB (in blue). Notably, the mean frequency of OM-SB was significantly more negative compared to POPC/PEG-PE SLB (n=3; p=0.0032). **(B)** Dissipation analysis of OM-SB (depicted in purple) and POPC/PEG-PE SLB (in blue). The mean dissipation of OM-SB was higher than that of POPC/PEG-PE SLB (n=3; p=0.0151). In both figures, 1. represents the injection of vesicles (OM-Hybrid or POPC/PEG-PE) into the flow cell, while 2. represents the injection of the wash buffer.

Based on the observed frequency shifts, we conclude that the OM-SBs are unilamellar. Previous work performed by Heath et al. introduced a method for the formation of the multi-layered SLB^62^. In their investigation, the frequency of a two-layer SLB typically doubles that of a unilamellar SLB. In contrast, OM-SBs exhibited a final frequency shift of -39.5 Hz, only 9 Hz lower than our POPC/PEG-PE bilayer. If the OM-SB were multilamellar, we would expect a double POPC/PEG-PE bilayer, resulting in a frequency shift of approximately -60 Hz. This observation strongly supports the notion that the OM-SB is indeed unilamellar.

### OMV-associated DNA is present on OM-SB

Our objective was to develop a surface that mimics bacterial outer membrane. Consequently, we conducted tests to detect the presence of bacterial OMV components within the surface. Our primary goal was to detect the presence of surface-associated DNA and proteins which are present on OMVs^63–66^. To accomplish the surface-associated DNA detection, we first created POPC/PEG-PE SLBs and OM-SBs labeled with the DiI membrane dye. Subsequently, we introduced YO-PRO-1, a cell-impermeable DNA-binding dye that fluoresces when bound to nucleic acids. YO-PRO-1 is frequently employed to monitor cell death, as it detects the exposure of cellular DNA during apoptosis^67,68^. Our observations revealed that the dye exhibited binding affinity exclusively to OM-SB, while it did not interact with the POPC/PEG-PE SLB surface **(Figure 6)**. This result establishes that OM-SB contains components characteristic of OMVs. Furthermore, this outcome underscores the gentle nature of our FT method, as the DNA associated with bacterial OMVs remained firmly attached to the surface throughout our experimentation. Moreover, this finding provides evidence for the exposure of the outer leaflet of the bacterial membrane since the dye is membrane impermeable therefore it cannot detect cytoplasmic DNA. This observation aligns with reports suggesting that DNA is situated on the exterior of these vesicles and is reinforced by the impermeability of YO-PRO-1 to the membrane^52,63,64^.

**Figure 6:**
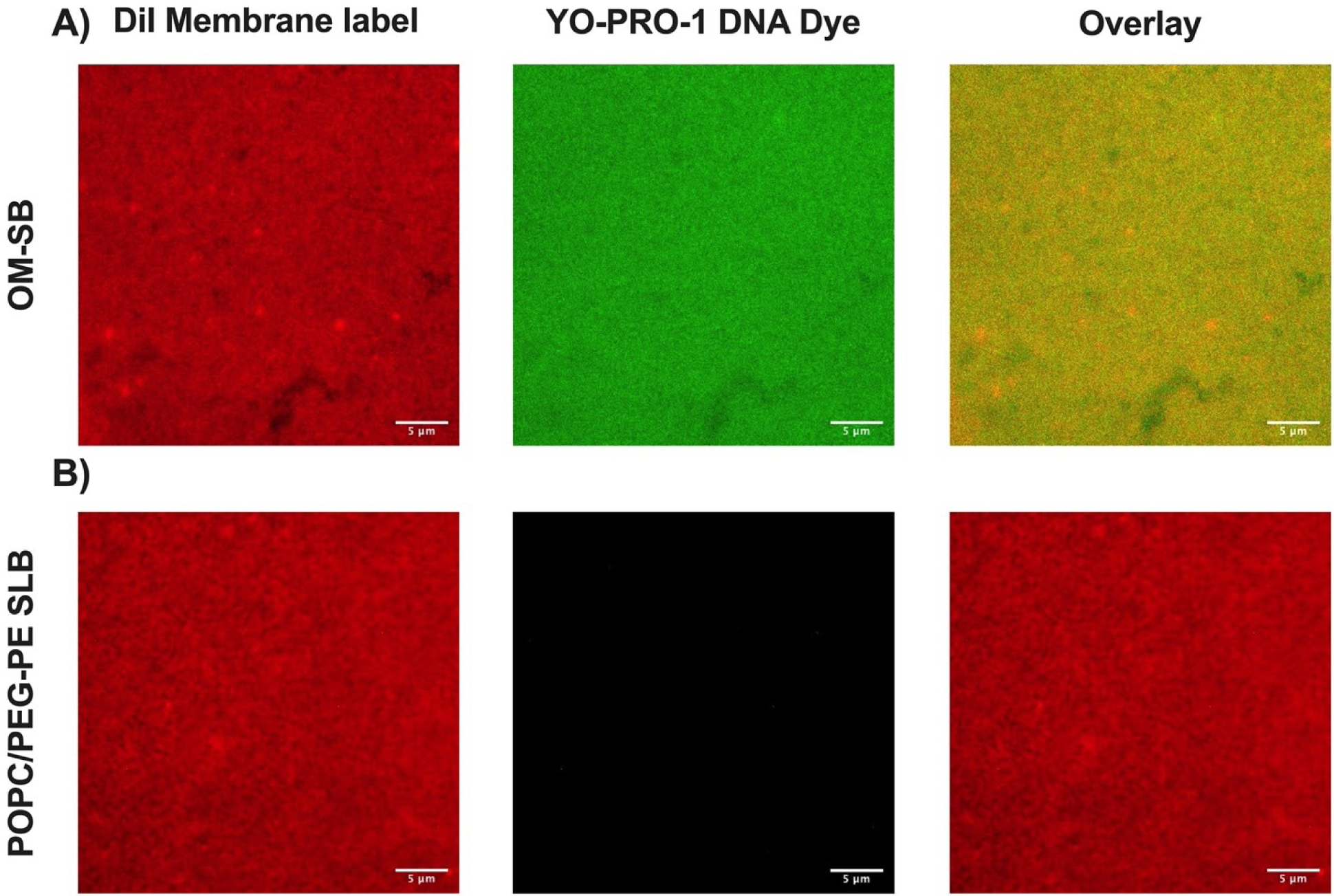
Detection of surface-associated DNA. (A) Fluorescence images of OM-SB. (B) Fluorescence images of POPC/PEG-PE SLB. In both cases, the left column represents the bilayers labeled with DiI membrane stain, the middle column depicts the YO-PRO-1 dye, and the right column is the overlay, indicating specific binding of the YO-PRO-1 to OM-SB. Scale bar in all images = 5 µm.

### Labeling proteins in OM-SB

In order to test for the presence of proteins, we used the NHS Atto 488 fluorophore^30^. This method enabled us to label primary amines present on proteins on the surface, as well as any PE lipids potentially existing in the bacterial membrane. We prepared two bilayers: one composed of POPC/PEG-PE synthetic lipid and another of OM-SB. After allowing the bilayers to react with the NHS Atto 488 and subsequent washes, we observed a significant increase in mean fluorescence intensity of Atto 488 bound to OM-SB compared to the POPC/PEG-PE bilayer **(Figure 7)**. This suggests that our platform preserves OMV-associated proteins and lipids, and it can be used for OMV protein detection.

**Figure 7:**
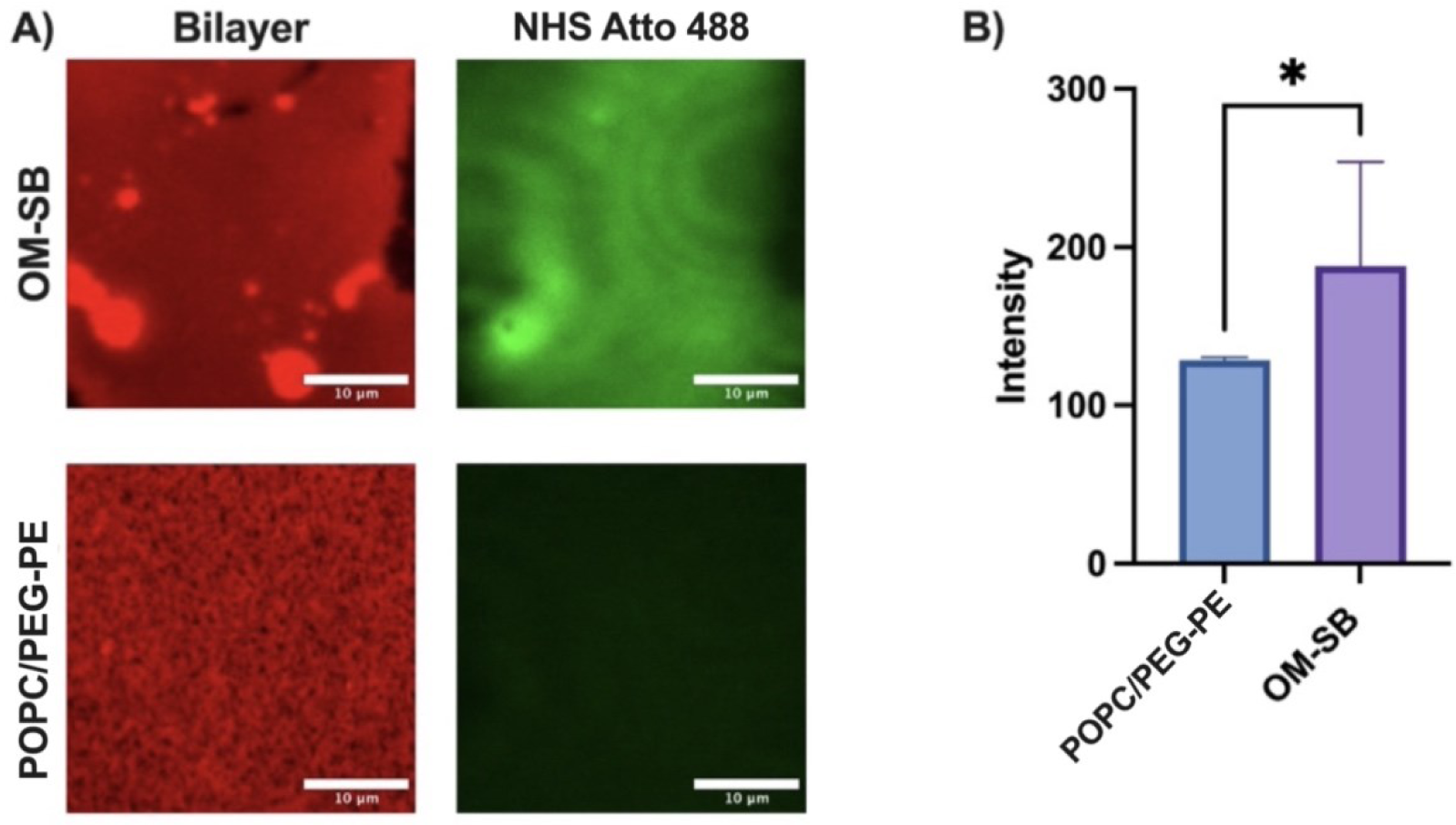
Detecting protein in OM-SB: A) micrographs of the bilayer and the NHS ester binding to OM-SB and POPC/PEG-PE. (B) Mean intensity profile of NHS Atto 488 (n=10).

### FITC-polymyxin B interaction with OM-SB

To further validate the effectiveness of our platform in studying antimicrobial peptide and OM-SB interactions, we focused on observing the interaction between polymyxin B and OM-SB. Previously, it has been reported that polymyxin B interacts with LPS that is present on the outer membrane of gram-negative bacteria, making it an ideal candidate for our platform^69,70^. OM-SB and POPC-PEG/PE bilayers were prepared as described previously and imaged **(Figure S12)**. FITC-labeled polymyxin B was then added to both bilayers, incubated for 1 hour, and subsequently imaged. For OM-SB, we observed significant polymyxin B aggregation, a phenomenon not seen in the POPC-PEG/PE bilayer **(Figure 8)**. Fluorescence intensity profiles revealed that in regions where peptide aggregation occurred, membrane defects and aggregation also occurred. It has been previously reported that polymyxin B inserts into the membrane, causing bacterial membrane disruption^69,70^. We hypothesize that as polymyxin B inserts into the membrane it begins to aggregate. This insertion and aggregation disrupts and aggregates the lipid bilayer structure. This demonstrates that our platform can be used to study antimicrobial peptide and membrane interaction successfully.

**Figure 8:**
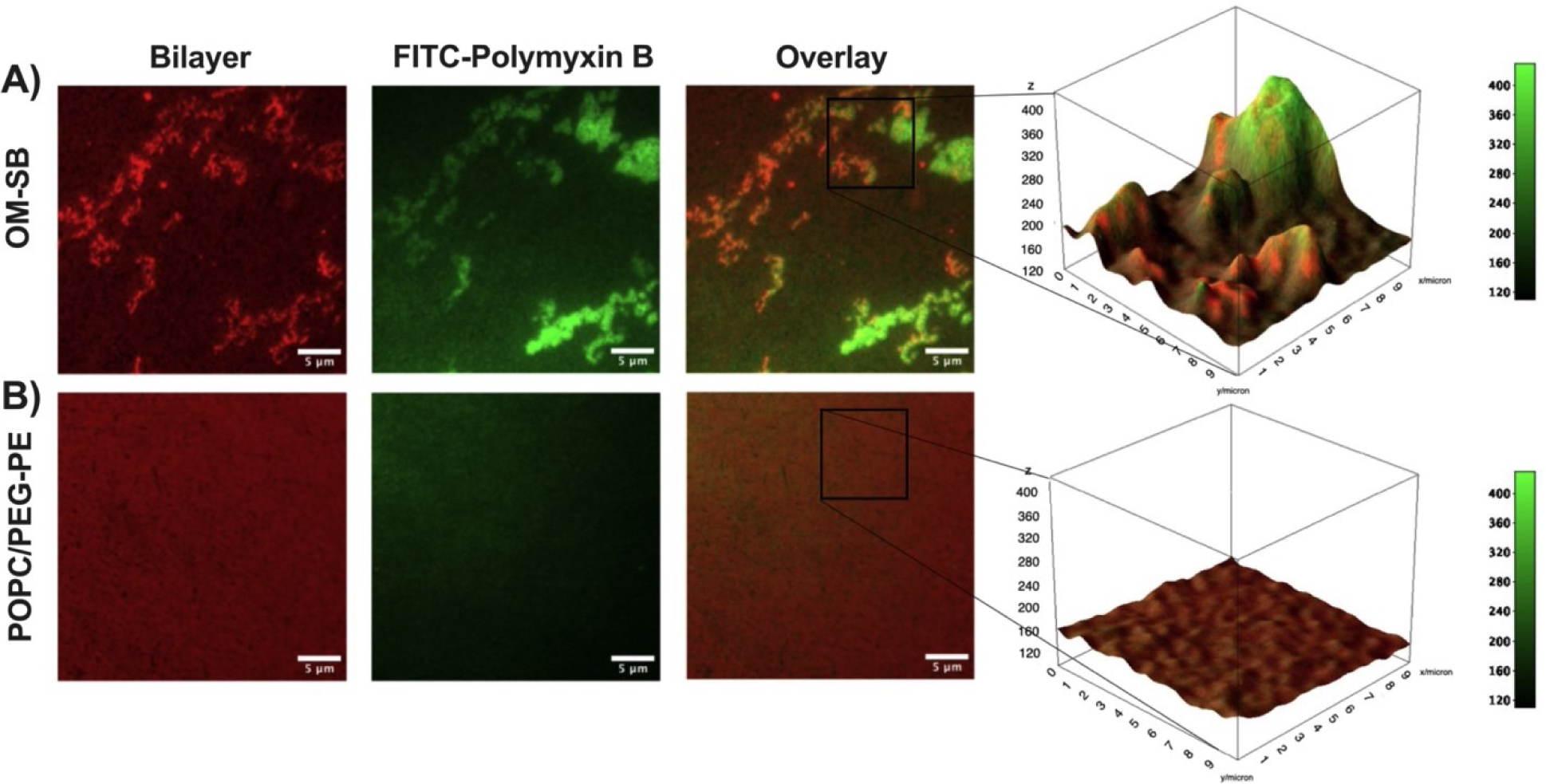
FITC-Polymyxin B and membrane interaction: **A)** OM-SB interaction with FITC-polymyxin B and **(B)** POPC-PEG/PE interaction with FITC-polymyxin B. The 3D intensity plot illustrated the FITC channel intensity (polymyxin B) overlaid on the TRITC channel (lipid bilayer).

## Conclusion

Amid the escalating concern surrounding antibiotic-resistant bacteria, there is an urgent demand for expediting drug development processes to effectively address this critical challenge. In this context, we present an innovative approach involving the creation of bacterial OM-Hybrid vesicles which spontaneously rupture on surfaces form a planar bacterial membrane that is compatible with surface-based biosensors, such as QCM-D. Our method demonstrates that incorporating 0.2 mg/mL of synthetic lipid with OMVs results in the formation of a planar surface that mimics the bacterial membrane. The resulting OM-SB membranes exhibit distinct lipid diffusion, and QCM-D frequency and dissipation properties compared to POPC/PEG-PE SLB controls. These variations strongly indicate the presence of biological materials (beyond phospholipids) in the OM-SBs. Intriguingly, our method also preserves non-covalently associated molecules, as we successfully labeled proteins and detected OMV-specific surface-associated DNA in the OM-SB. This emphasizes the significance of our platform in unraveling bacterial physiology, as surface-associated DNA plays pivotal roles in various critical physiological processes, including biofilm formation which aids in antibiotic resistance^71^. This discovery reinforces the effectiveness and reliability of our approach in exploring planar surfaces generated from bacterial OMVs.

Daniel and colleagues have previously introduced a method that facilitates the creation of a planar surface through bacterial OMVs. While this approach offers certain advantages, it also presents a few drawbacks, such as the absence of a direct measure for determining the extent of POPC/PEGPE interaction with OMVs and the resulting surface coverage. Moreover, this method necessitates a higher concentration of synthetic lipids. On the other hand, one advantage of this approach compared to ours is the absence of an additional step required for OM-Hybrid formation. In contrast, our method not only enables the generation of these hybrids which may be applicable to drug delivery and antibiotic loading. Previously, amphipathic peptides have been employed to form synthetic lipid bilayers on non-conventional surfaces such as gold^72^. While there are reports of EV and OMV binding peptides, their use to form a bilayer surface remains limited^73,74^. Our platform holds considerable significance in the realm of pharmaceutical and microbiological research. It offers an authentic mimic of the bacterial surface environment, providing an ideal substrate for testing and developing new antibacterial agents.

## Supporting Information

Fluorescence micrographs, FRAP recovery curves, fluorescence spectra, zeta potential measurements, and QCM-D data (PDF). Fluorescence microscopy movie of OM-SB formation (AVI).

## Acknowledgements

This work was supported by grants from the National Institutes of Health (R21GM134414, R21DE032153, and R15GM152918), and the Pennsylvania Department of Health (4100095607). We gratefully acknowledge Anand Ramamurthy for providing access to the Zetasizer. Additionally, we extend our appreciation to Marcos Pires for supplying the FITC-labeled polymyxin B.

## For table of contents only

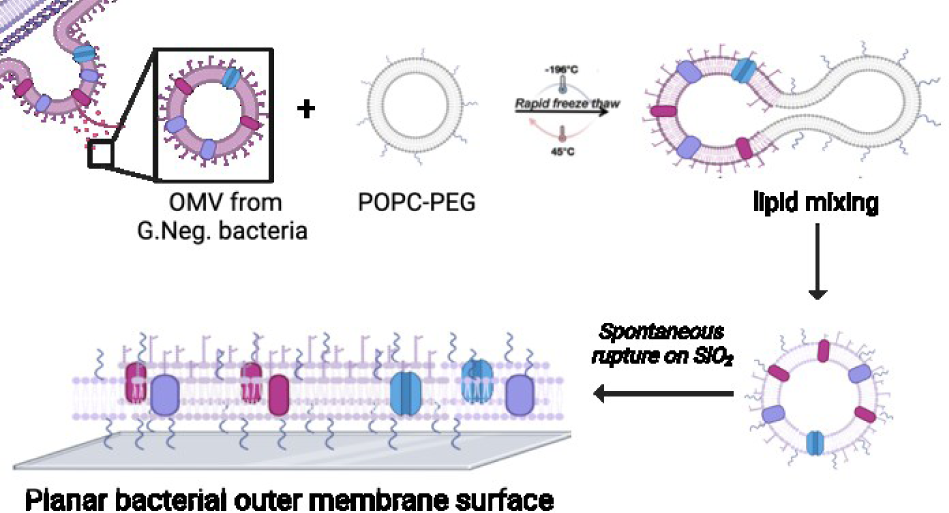

