## Supplemental Information for "Engineering Planar Gram-Negative Outer Membrane Mimics Using Bacterial Outer Membrane Vesicles"

**Supporting Information**

Aarshi N. Singh[1], Meishan Wu[2], Tiffany T. Ye[1], Angela C. Brown[2], Nathan J. Wittenberg[1*]

[1]Department of Chemistry, Lehigh University, Bethlehem, PA, USA
[2]Department of Chemical and Biomolecular Engineering, Lehigh University, Bethlehem, PA, USA

*Corresponding author: Nathan J. Wittenberg


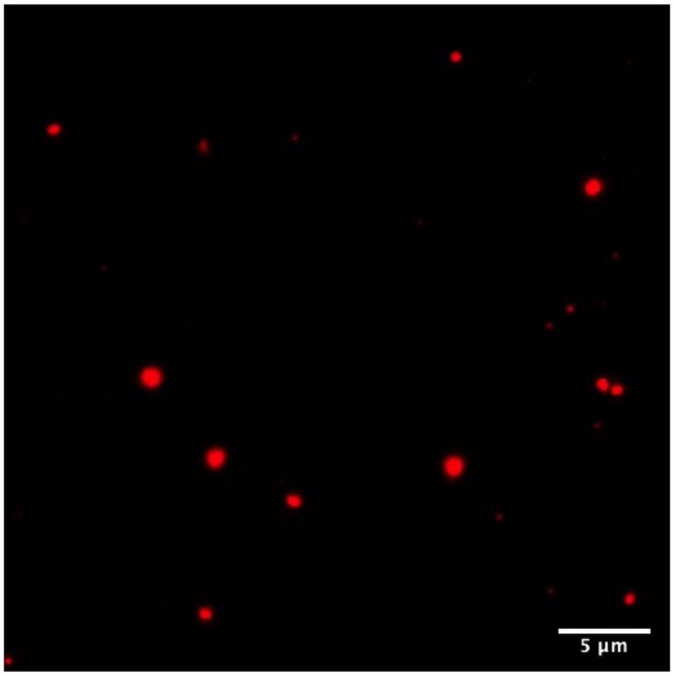


**Figure S1:** ***OMVs adsorb but do not rupture on glass***. Native OMVs were added to the clean glass and OMV accumulation was observed, where they did not rupture to form a planar surface. Scale bar in all images = 5 µm.

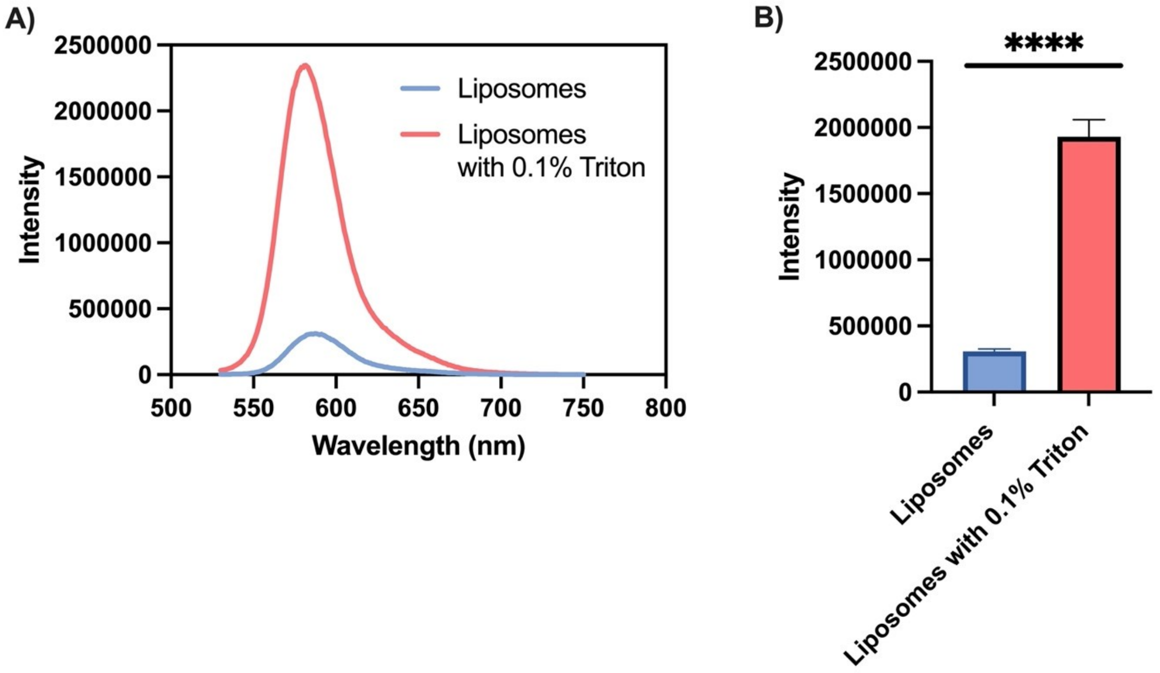


**Figure S2: (A)** Fluorescence emission spectra of liposomes with 10 mole % R18 and liposomes with 0.1% Triton to induced lysis. **(B)** A significant increase in fluorescent intensity ($P < 0.0001$; $n = 3$) was observed following the lysis of liposomes using 0.1% Triton.

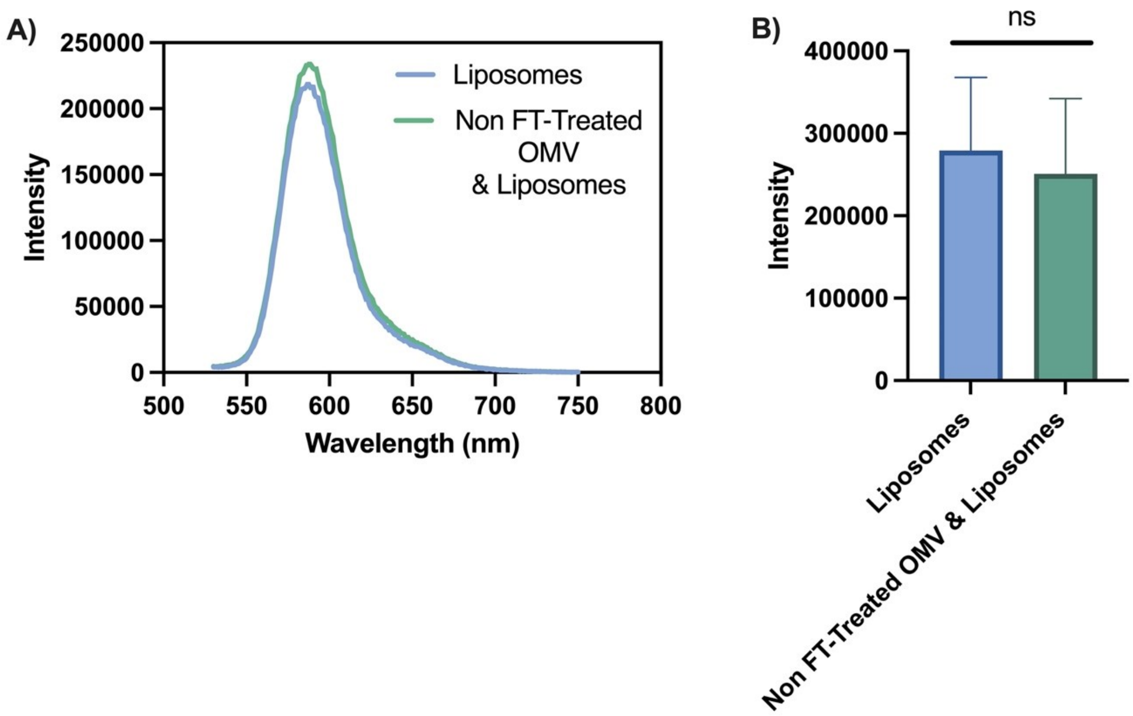


**Figure S3 (A)** Representative fluorescence emission spectra of liposomes and non FT-treated liposomes and OMVs. **(B)** A non-significant increase in fluorescent intensity (n = 5) was observed indicating that FT-treatment is required for lipid mixing.

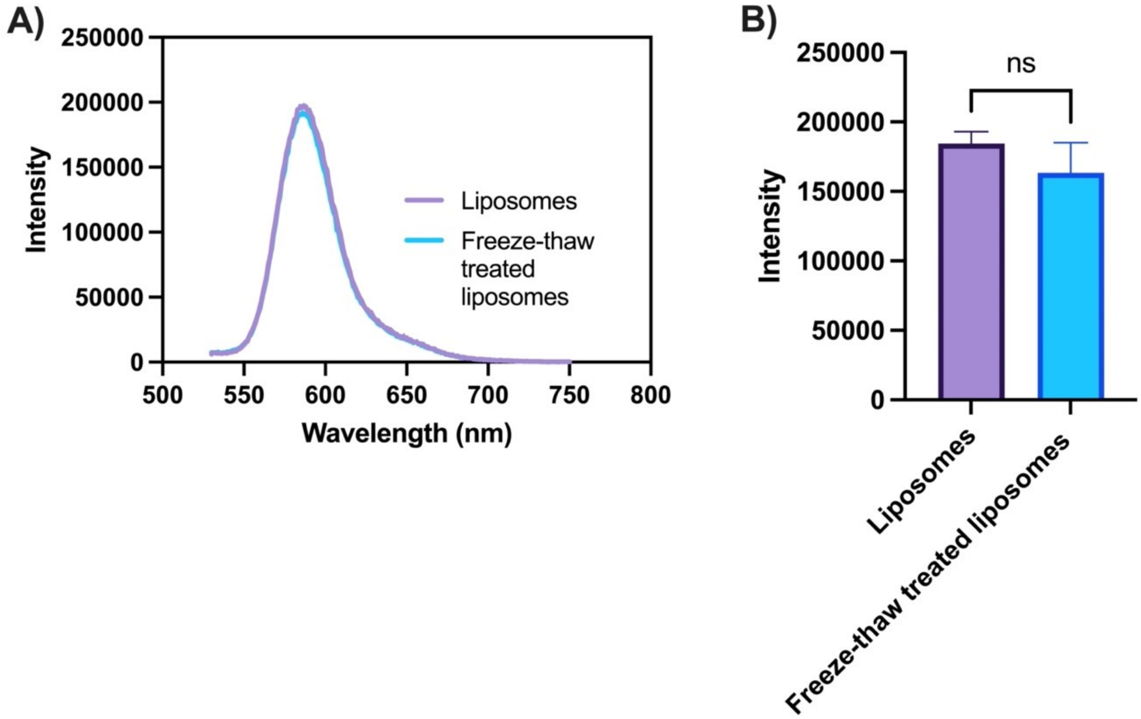


**Figure S4** ***R18 liposome freeze-thaw treatment control:*** **(A)** Representative spectra of R18 liposomes vs freeze-thaw treated R18 liposomes**. (B)** A non-significant change in the intensity was observed between R18 liposomes and freeze-thaw treated R18 liposomes (n=3).

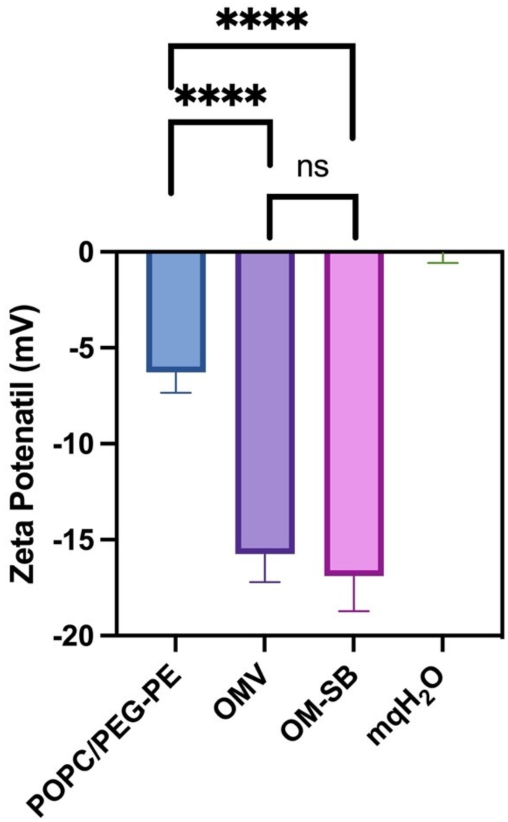


**Figure S5** ***Zeta potential measurement:*** The average zeta potential is -6.278 ± 1.057, -15.74 ±1.47, -16.89 ± 1.83, -1.090 ± -0.064 of POPC/PEG-PE, OMV, OM-SB, and mqH2O, respectively.

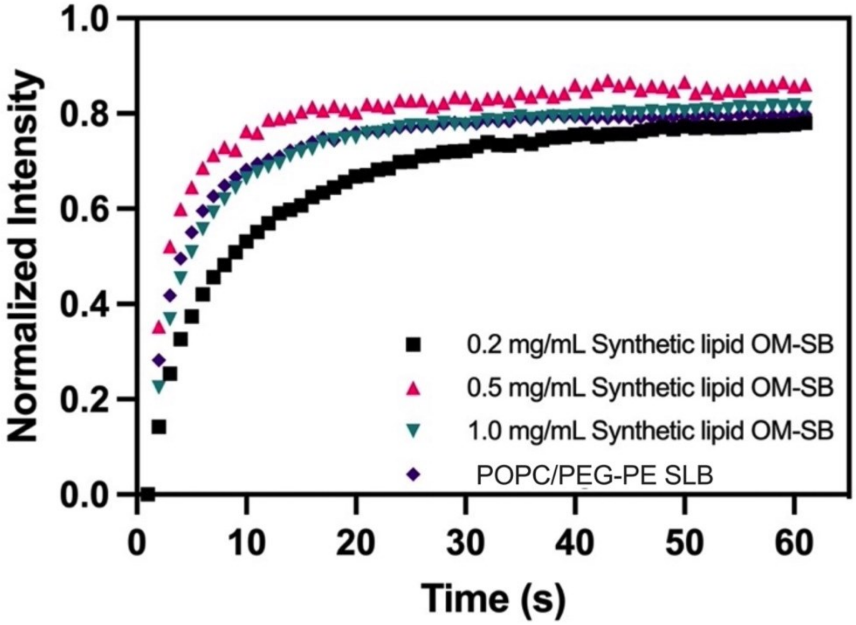


**Figure S6:** ***FRAP curves of varying synthetic lipid component in OM-SB***: FRAP curves were generated using varying concentrations of POPC-PEG in OM-SB formation, with fluorescence recovery after photobleaching recorded. Average curves shown based on n≥3 measurements.

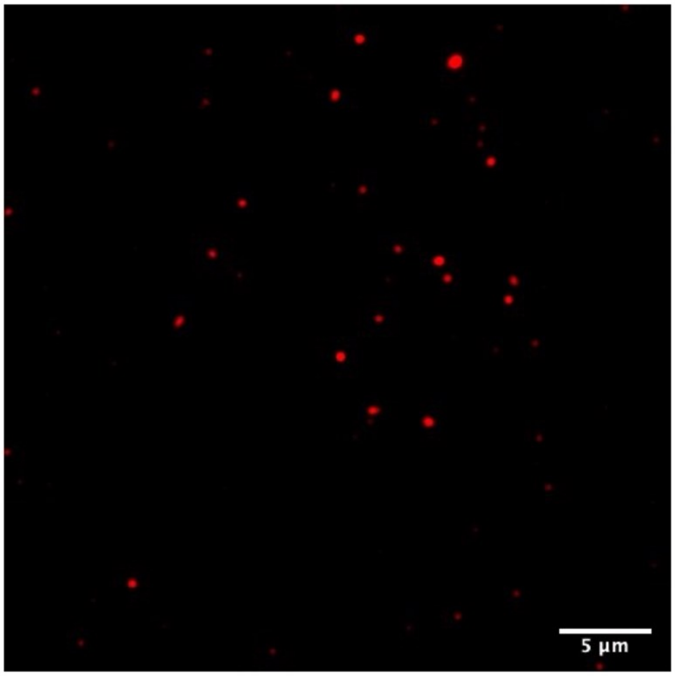


**Figure S7:** ***FT-treated OMVs do not rupture on glass:*** Intact FT-treated OMVs were observed on the clean glass surface indicating that FT treatment alone is not sufficient to induce OMV rupture. Scale bar in all images = 5 μm.

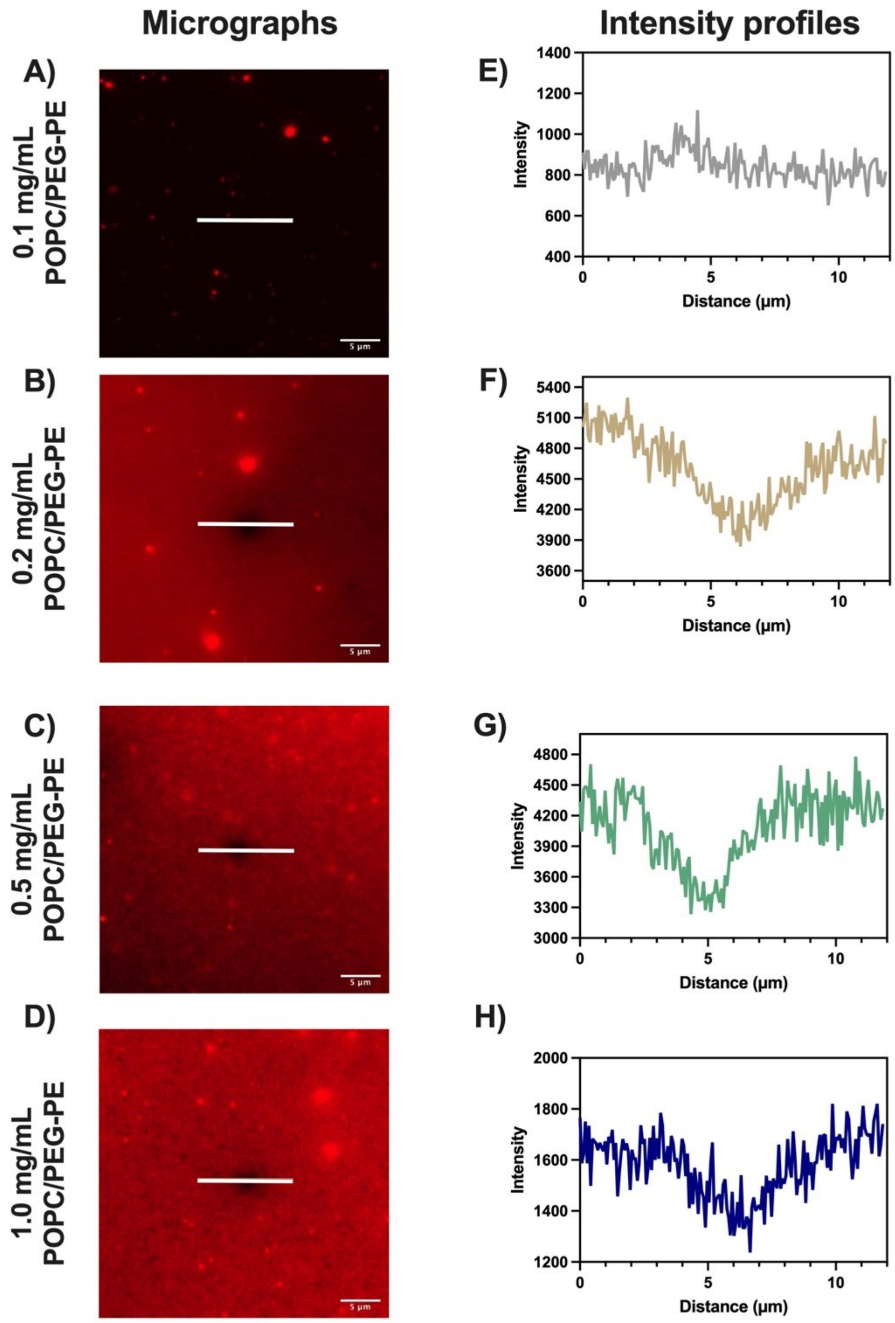


**Figure S8:** ***Effect of different concentrations of POPC-PEG used in OM-Hybrid formation:*** **(A-D)** micrographs exhibit varying concentrations of synthetic lipids, specifically, **(A,)** 0.1 mg/mL POPC-PEG **(B)** 0.2 mg/mL POPC-PEG **(C)** 0.5 mg/mL POPC-PEG **(D)** 1.0 mg/mL POPC-PEG. The white line denotes the scanned area for intensity profiling, represented in figure **(E-H).** Scale bar in all images = 5 µm.

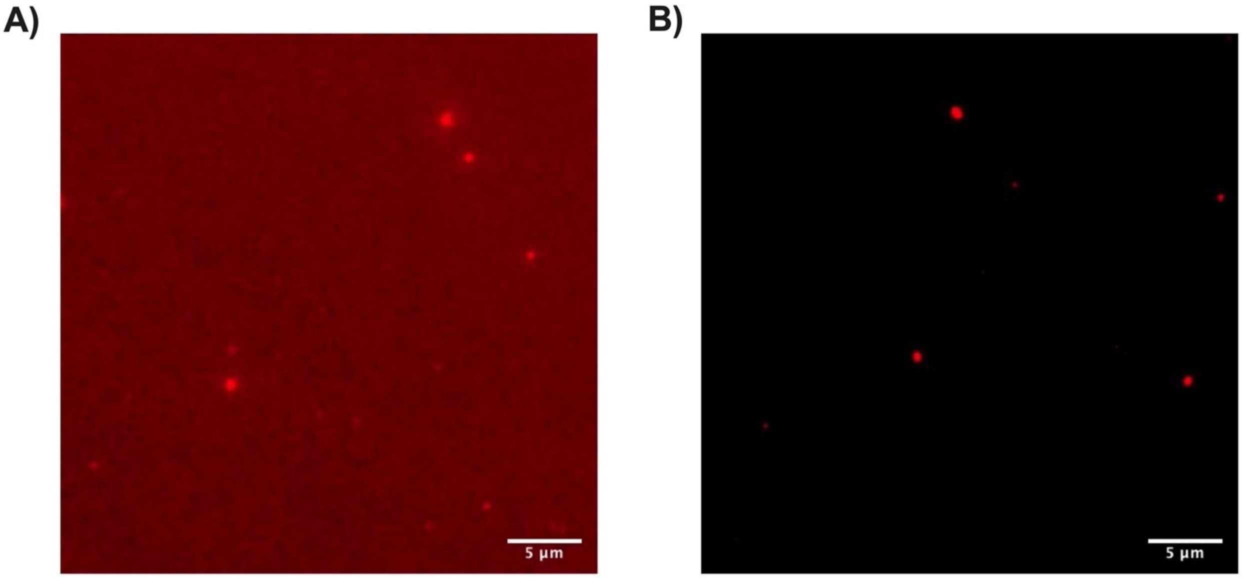


**Figure S9:** ***FT-Treatment of OMV and liposomes is required to form a bilayer:*** **(A)** Fluorescence image of FT-treated OMV-Liposome mixture (OM-Hybrids) exposed to bare glass. **(B)** Fluorescence image of non FT-treated OMV-liposomes mixture exposed to bare glass. Scale bar in all images = 5 μm.

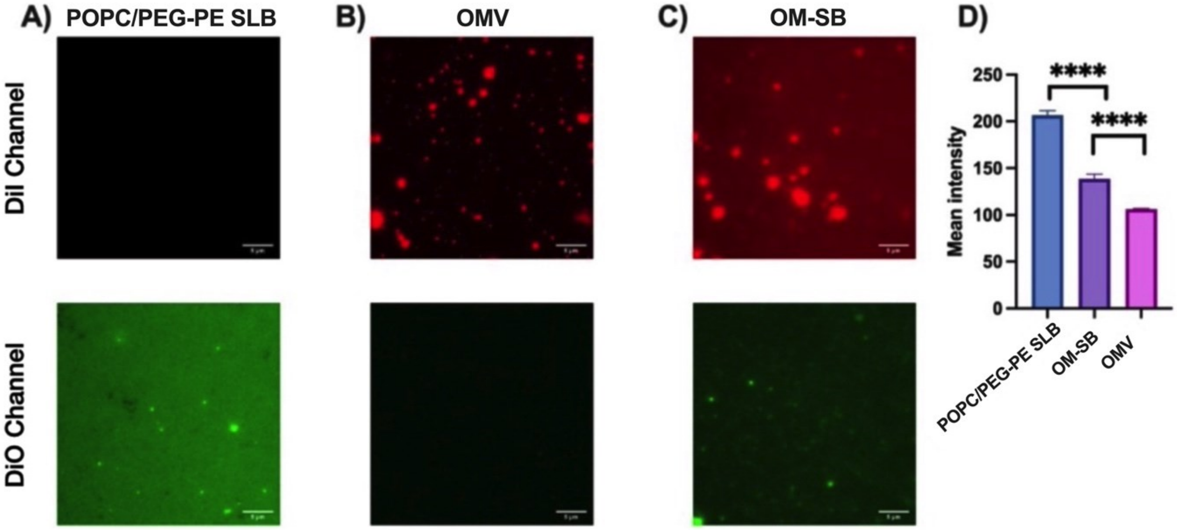


**Figure S10:** ***Analyzing the POPC/PEG-PE component in OM-SB*** **(A)** The POPC/PEG-SLB, prepared with DiO-labeled liposomes, was utilized at a concentration of 0.2 mg/mL to mimic the concentration in added to prepare OM-SB. **(B)** OMVs labeled with DiI added to a clean glass at 4.5 mM lipid concentration. **(C)** OM-SB prepared with 4.5 mM lipid concent of DiI labeled OMVs and the 0.2 mg/mL of DiO labeled POPC/PEG-PE liposomes. Both fluororophores were observed in the bilayer indicating a success incorporation both of the componenets. **(D)** Quantification of DiO fluorescence in both POPC/PEG-PE SLB and OM-SB revealed a significant reduction in fluorescence intensity ($p < 0.0001$), indicating that only 32% of OM-SB is composed of POPC/PEG-PE.

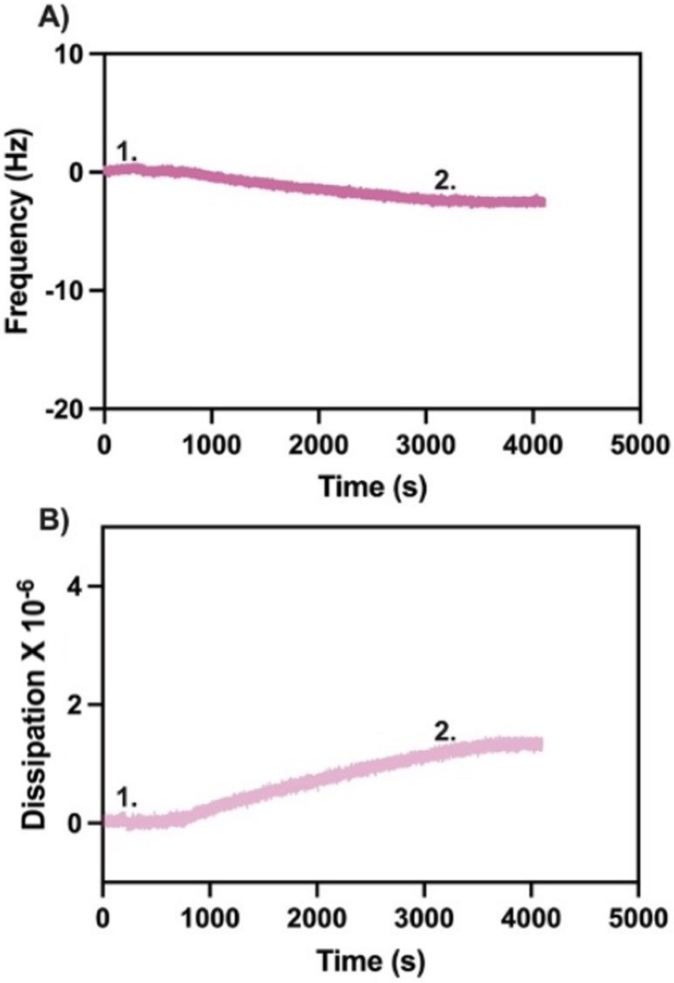


**Figure S11:** ***Frequency and dissipation assessment of OMV adsorption on a QCM-D Sensor*: (A)** Frequency shift of OMVs adsorbing to a QCM-D Sensor. The observed frequency shift due to OMV adsorption to the QCM sensor was -2.51 Hz. **(B)** Dissipation shift of OMVs adsorbed to QCM-D Sensor. The observed dissipation associated with OMV adsorption to the QCM was 1.33 ppm. In both figures, 1. Represents the injection of OMVs into the flow cell, while 2. represents the injection of the buffer.

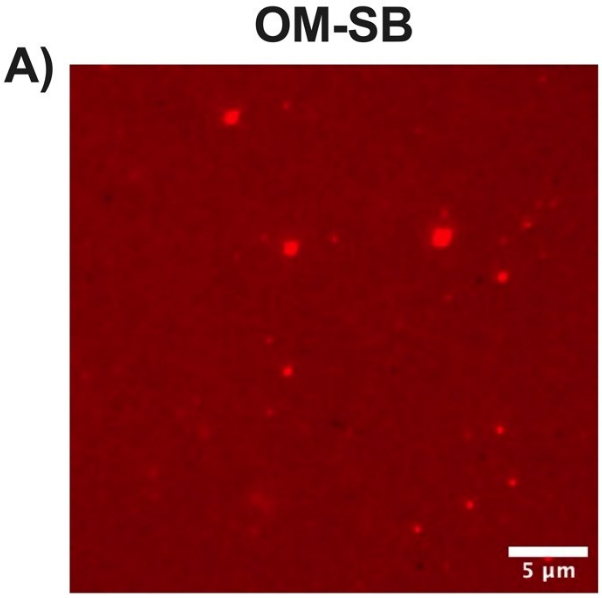


**Figure S12:** ***Micrographs of bilayers before treatment with Polymyxin B:*** **(A)** OM-SB and **(B)** POPC-PEG/PE
